# Critical pollination chemistry: Specific sesquiterpene floral volatiles in carrot inhibit honey bee feeding

**DOI:** 10.1101/2022.10.11.511710

**Authors:** Stephen R. Quarrell, Alyssa M. Weinstein, Lea Hannah, Nicole Bonavia, Oscar del Borrello, Gavin R. Flematti, Björn Bohman

## Abstract

- Although many plant species are reliant on insect pollination, agricultural plant breeding programs have primarily focused on traits that appeal to growers and consumers, rather than on floral traits that enhance pollinator attraction. In some vegetable seed production systems, this has led to declining pollinator attraction and poor seed yields.
- We predicted that low-yielding crop varieties would be less attractive to pollinators due to deficiencies in nectar rewards or volatile floral attractants. To test our prediction, we used a chemical phenotyping approach to examine how floral chemical traits of five carrot lines affect honey bee visitation.
- In bioassays, honey bees avoided feeders containing nectar from all carrot lines indicating a general non-attractant effect. Certain compounds in carrot flowers and nectar not only failed to elicit attraction but functioned as repellents, including the sesquiterpenes α-selinene and β-selinene. Others enhanced attraction, e.g. β-ocimene.
- The repellent sesquiterpenes have previously been implicated in plant defense suggesting a fine balance between pollination and plant protection, which when disrupted in artificial selection in plant breeding programs can impact the crop yield. These new insights highlight the importance of bioactive compounds in attracting pollinators toward floral resources in both ecological and agricultural settings.

## Introduction

Ecosystem function is underpinned by numerous biotic and abiotic factors. In angiosperms, effective pollination is crucial for sexual reproduction and in many instances is facilitated by insect pollinators (Ollerton *et al*., 2011). When foraging, pollinators typically show floral preferences that are mediated by a suite of visual, olfactory, and gustatory cues (Dötterl & Vereecken, 2010). The key traits that attract pollinators are considered to include direct rewards such as nectar (Beekman, 2005; Nepi, 2017) and pollen (Arenas & Farina, 2012; Stabler *et al*., 2018), attractive floral volatiles, and visual cues (Dötterl *et al*., 2014; Parachnowitsch & Manson, 2015). However, cross kingdom interactions can be complex as social insects do not solely use direct cues of attraction when foraging, and may heed or ignore resources based on a variety of other associative factors communicated by siblings. These decision making processes may include other stimuli including the presence of predators (Avarguès-Weber *et al*., 2018), shifts in the prevalence of a specific resource, or potentially the perceived quality of a specific resource relative to others that become available within the foraging range (Quinlan *et al*., 2021). For these reasons, physical attributes of flowers alone do not necessarily trigger foraging behaviour instantly and insect behaviour can change once a particular cue has been encountered (Kevan & Baker, 1983). Many pollinators display flexibility in these preferences due to associative learning between rewards and floral characteristics (Schiestl & Johnson, 2013). The interaction between these cues can shift floral visitation from abundant resources toward higher quality, less abundant forage. For example, many agricultural crops that have a periodic over-abundance of floral resources may not always possess the nectar rewards or attractive floral cues required to induce pollinator visitation (Gaffney *et al*., 2011; Topitzhofer *et al*., 2019). Therefore, it is essential to consider insect learning processes to understand pollination efficiency, particularly within agricultural production systems (Jones & Agrawal, 2017).

With over 35% of global food crops at least in part dependent on animal pollination (Klein *et al*., 2007; Potts *et al*., 2016), the foraging preferences of pollinators within agricultural crops is being increasingly recognised as an important factor in maintaining agricultural production levels and quality (Reilly *et al*., 2020). Economically, it has been estimated that nearly 10% of the total value of agricultural production, or ∼US$200B is derived by insect pollination (Hristov *et al*., 2020).

Although many animals are important pollinators, the European honey bee *Apis mellifera* remains the most widely used pollinator for commercial crops (Klein *et al*., 2007). However, diseases, parasites such as the Varroa mite, use of pesticides, and other factors have led to a drastic decline in honey bee numbers in recent decades (van Engelsdorp *et al*., 2009; Barbosa *et al*., 2015; Kang *et al*., 2016). Strategies to safeguard global food production in the face of drastic declines in honey bee numbers include an enhanced focus on alternative pollinators (Garibaldi *et al*., 2013; Garibalidi *et al*., 2016) and revised crop management strategies (van Gils *et al*., 2016; Garibaldi *et al*., 2018). No doubt the question of securing pollination services, both for biological conservation and food production, deserves a broad focus. Nonetheless, there is also an immediate knowledge gap regarding the importance of chemical cues with respect to pollinator floral preferences to agricultural crops (Stevenson *et al*., 2017). Firstly, despite the common use of managed honey bee hives in agricultural production systems, pollen transfer between flowers often remains limited, especially in crops that do not naturally depend on bees for pollination (Westerkamp, 1991; Lamborn & Ollerton, 2000; Mas *et al*., 2018). By increasing hive density, crop pollination rates and subsequent seed set can be improved, but for some varieties, such measures remain insufficient with low yields still common (Gaffney *et al*., 2019). Secondly, several crops grown for human consumption are selectively bred for traits favoured by growers and consumers, such as elevated pest and disease resistance and appealing taste and physical appearance, but are seldom bred for pollinator attraction (Nothnagel & Straka, 2000). This trend has led to pollination deficits becoming increasingly noticeable, particularly with the introduction of hybrid crop varieties. Several modern crops experience reduced seed yields despite managed honey bee hives being used in excess (Aizen *et al*., 2008).

Honey bees are well known to associate rewards with phenotypic cues such as colour and floral volatiles (Jones & Agrawal, 2017). The influence of colour and floral morphology in particular has been studied in detail for bee pollination (*e*.*g*. Menzel, 1968; Giurfa *et al*., 1995). Yet, despite their importance particularly within vegetable seed production, the detailed cues underpinning honey bee attraction to vegetable flowers remain poorly understood. In flowering hybrid crops, bees may initially orientate toward visual cues in flowers of one cultivar or accession. Once these flowers are located, different more attractive chemical cues from different neighbouring cultivars or accessions could override the initial attractive signals. It has been established that although possible, it is difficult to train social insects by manipulating associative cues without linking these traits with a reward (Seeley *et al*., 1991; Menzel & Müller, 1996; Beekman, 2005). What appears to be much less known is how and why certain floral traits, which appear not to be linked with rewards, are avoided. In such cases, for example olfactory cues, where the lack of attraction appears to be independent of the degree of reward, there may be scope to affect the insect behaviour by manipulating or reducing the trait, thereby improving pollinator attraction. The accepted importance of floral volatiles, however, has led to the development of pollination-improving methodologies relying on the conditioning of pollinators to odours that are abundant in the flowers of target crops. By manipulating the pollinators, often honey bees, to associate these odours with rewards, the yields can be improved, but not always (Silva *et al*., 2003). Apart from the costly and time-consuming process involved in pollinator pre-conditioning, the long-term efficiency of these methodologies is questionable, as the insects learn that the level of reward is not maintained (Silva *et al*., 2003).

Pollinator attraction is often measured by quantifying pollination rates and improved seed yields (Rachersberger *et al*., 2019; Seo *et al*., 2019; Mas *et al*., 2020). To decipher chemical attraction, detection of pollinator responses to individual selected chemical components of nectar or the floral headspace can be measured with antennal electrophysiology methods or honey bee proboscis extension response bioassays (Henning & Teuber, 1992; Liu *et al*., 2015; Twidle *et al*., 2015; Barazani *et al*., 2019). However, routine electrophysiology methods cannot detect all biologically active compounds, nor distinguish between the different functions of the semiochemicals being assessed (*e*.*g*. if attractant, repellent, or other) (Bohman *et al*., 2016).

Proboscis extension response bioassays suffer from a strong association between the response and reward, or in other words, responses are linked to memory effects (Pham-Delegue *et al*., 1993), with any compounds that the bees have not been conditioned to potentially going undetected. Consequently, to observe the natural behaviour of the pollinators, full behavioural bioassays are preferred (Schreiner *et al*., 2013). The importance of specific volatiles from crop flowers in honey bee attraction has previously been demonstrated in field bioassays with artificial flowers and alfalfa (Henning *et al*., 1992).

Carrots (*Daucus carota* L.) are an important commercial crop, for which it has been established that pollination limitation is a major contributor to low seed yields (Spurr, 2003) and multiple studies have shown that the pollination rates of hybrid varieties are lower than in open-pollinated (OP) varieties (Erickson *et al*., 1979; Galuszka & Tegrek, 1989). Compared to alternative pollinators (Davidson *et al*., 2010; Howlett, 2012; Howlett *et al*., 2017; Gaffney *et al*., 2018), as for most crops, honey bees remain the most effective pollinators of hybrid carrot (Gaffney *et al*., 2018). Thus, hybrid carrot represents a suitable model for investigating variables that negatively affect pollination success in honey bee pollinated cropping systems. The effects of colour and floral morphology have been studied for bee pollination of carrot, with inflorescence color and nectar sugar composition and concentration found not to differ significantly between hybrid lines (Gracie, 2011; Gaffney *et al*., 2020). However, no comprehensive studies of the role of volatiles have been reported. In this study, we compare the attractiveness of four parental carrot accessions including two sterile, pollen free, Cytoplasmic Male Sterile (CMS) carrot lines and their reciprocal fertile pollen bearing, fertile ‘maintainer’ lines. Each pair bears the same nuclear genome with the sterile CMS parent being emasculated via the genetic manipulation of the *restorer of fertility* (R*f*) genes within the plant’s cytoplasm (Chen & Liu, 2014). Industry documented seed set data for each pairing indicated each reciprocal pair (fertile and sterile) yielded either consistently low or moderate seed yields. These accessions were compared to a fifth open-pollinated (OP) cultivar, Western Red. The OP cultivar is known to produce high seed yields and is considered highly attractive to honey bees. We hypothesised that nectar from the carrot accessions producing less seed would be less attractive to honey bees and aimed to determine volatile compounds contributing to differences in honey bee attraction to different carrot accessions. The traits of nectar composition (floral reward; sugar and micronutrient concentration and composition) and volatile composition (floral attraction) were investigated. Bee attraction to individual characteristic chemical compounds of specific accessions was tested and confirmed in behavioural bioassays.

## Materials and Methods

### Carrot lines

For the ongoing production of hybrid carrot seed, three parental lines are required: The A-line, (cytoplasmic male-sterile line used as the female parent;), the B-line is the ‘maintainer’ line (male parent to produce more female A-line) and a third R-line (restorer) which is the pollinator or fertile male parent, which is crossed with the A-line to produce hybrid seed (Chen & Liu, 2014). Carrot seed from the two reciprocal pairs of petaloid, parental male sterile (LS = low yielding or MS = medium yielding) and male fertile (LF = low yielding or MF = medium yielding) carrot accessions (Figure 1A) were supplied by seed company Rijk Zwaan Australia Pty Ltd. Industry documented seed set data for each pairing indicated each reciprocal pair (fertile and sterile) yielded either consistently low or moderate seed yields. Carrot seed from a fifth, high yielding, open-pollinated (OP) cultivar, Western Red (H = high yielding), was purchased from commercial seed supplier “The Diggers Club” (https://www.diggers.com.au/). All seed was planted in 15 cm plastic plant pots using commercially sourced potting mix and housed in a glasshouse at 23° C. Once fully grown, the carrot stecklings were vernalised to induce flowering by removing them from the pots, with any excess soil removed by washing in tap water. The stecklings were then dipped in the fungicide Mancozeb Plus (Yates, Product code: 53850) and buried in polystyrene boxes filled with moist river sand and refrigerated at 4 °C for 10 weeks. Once vernalised the carrots were repotted into 20 cm pots using the same commercially available potting mix and grown outdoors to flowering. For all four parental lines and the OP cultivar, umbellets were picked from the first, second and third umbel, between 10 am – 2 pm, 3-6 days after the opening of the first flower of each umbel. Samples were processed within one hour of collection.

**Figure 1.**
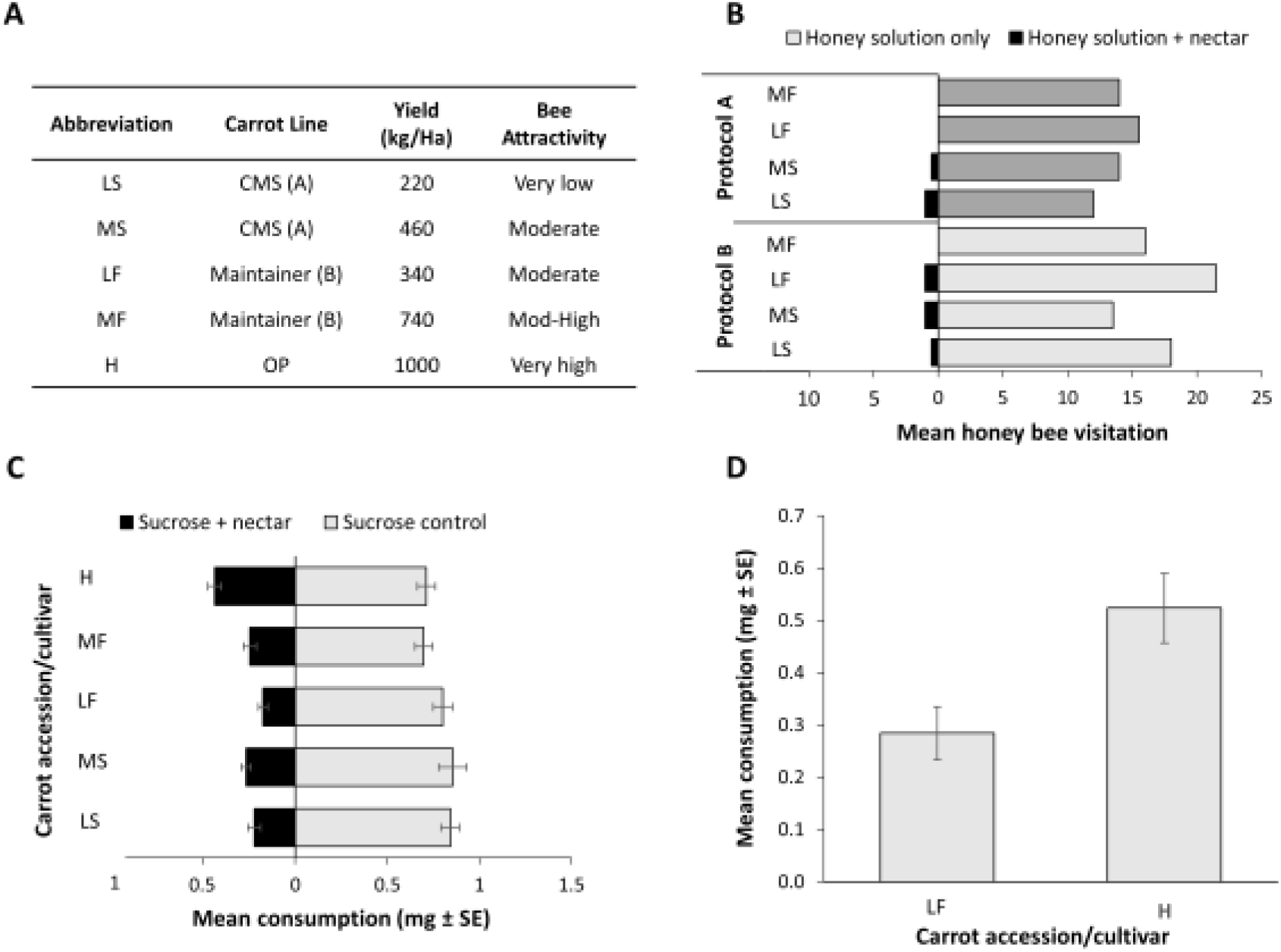
A Descriptions of four carrot accessions (LS, MS, LF and MF) and one open-pollinated cultivar, Western Red (H) used in this study, with seed yields and relative honey bee attractivity. B: Results from preliminary field bioassays. Number of bees feeding from each feeder. Controls consisted of honey solution, treatments of honey solution with added carrot nectar from each carrot parental accession (L and M) extracted with protocol A (dark grey bars) and protocol B (light grey bars). Controls were conducted pairwise with each treatment, with corresponding coloured bars. C: Results from laboratory bioassays, pairwise choice experiments comparing sucrose solutions with sucrose solutions spiked with carrot nectars for each accession and cultivar. D: Results from laboratory bioassays, pairwise choice experiment comparing sucrose solution spiked with nectar from line LF with sucrose solution spiked with nectar from variety H.

### Nectar extraction for bioassays and chemical analysis

For bioassays, carrot nectar was extracted following one of the following protocols. A) Spinning in Eppendorf tubes, in a modified method from Giralamo (1973). A spin filter (UltraClean Mini Plasmid Prep, Mo Bio Laboratories Inc, USA) in an Eppendorf tube was fully packed with individually collected carrot umbellets, with all stalks removed. Each sample was spun at 12000 rpm for 10 min at room temperature. The carrot tissue was removed, the filter re-packed with fresh umbellets and re-spun. The procedure was repeated until ca 200 µL of nectar was separated. B) To rule out any risk of contamination by co-extracted sap, not naturally accessible to the pollinators, method B, an alternative method without mechanical extraction, was also applied. In this semi-quantitative method, umbellets (n = 60) were individually dipped in water (2 mL), twenty times per umbellet, forming an extract of ca 200 µL. Nectar samples from both methods were stored at -20 °C until used in bioassays. For chemical analysis, nectar was extracted by protocol B with minor modifications: Umbellets (n = 20) were dipped, one-by-one, ten times in distilled water (1.0 mL). To the aqueous extract (ca 100 µL) in an Eppendorf tube, was added dichloromethane (100 µL). The sample was vortexed (10 s), and an aliquot (50 µL) of the organic layer was removed for analysis of volatiles. The remaining sample was concentrated under nitrogen and taken up in distilled water (100 µL). An aliquot (10 µL) was removed for carbohydrate analysis while the reminder was used for AES analysis.

### Field Bioassays

The bioassay setup consisted of feeders made from clear plastic specimen jars of 5 cm diameter, with blue lids on which Eppendorf lids were glued upside down. The treatments were added to these Eppendorf lids, which acted as dispensers. The feeders were placed on a fence ca 1 m from the ground, intercepting a grassy slope approximately perpendicular and 20 m away from two managed beehives (ca 30.000 bees per hive) at Sandy Bay, Tasmania. First, we confirmed that pure carrot nectar was not attractive to the honey bees within 4 hours, even after several attempts to train the honey bees to the feeders from nearby hives. Then, we conducted sequential experiments with volumes of 50 µL per test, in the following order: 1. Honey solution (20% sugar); 2. Treatment (Nectar extracted with method A or B, above); 3. Honey solution (20% sugar); 4. Treatment spiked with honey (3:1 treatment to honey solution); 5. Honey solution (20% sugar). All experiments were conducted in duplicates, on different days. As a control experiment, honey solutions were diluted four times, to exclude the variable sugar content as the cause of lack of feeding in the treatments. Diluted honey (5% sugar) still promoted feeding.

### Laboratory bioassays

All laboratory bioassays were conducted in a controlled temperature room at the University of Tasmania. The room conditions were 30-34 °C, with >50% humidity and an 8L:16D lighting regime. Bug Dorm (MegaView Science, Taiwan) cages, 25 × 25 × 25 cm were used with 10 foraging honeybees housed in each cage. Nectar foraging bees were collected from the hive entrances at the University of Tasmania’s research apiary, Sandy Bay, Tasmania. All bees were collected while leaving the hive and transported to the controlled temperature room within 30 min of collection. Bees from each cage were sourced from one hive only. Upon arrival, each cage was provided with 40% w/w sucrose solution from a feeder consisting of two Eppendorf tubes (2 mL) suspended ca 0.5 cm from the cage floor in an Eppendorf tube holder with the tubes spaced 10 cm apart. Each Eppendorf tube had 3 × 1.5 mm holes drilled into the terminal end to allow bee feeding. All experiments commenced upon arrival during the room’s photophase and lasted 48 hours. The first 24 hours allowed the bees to both acclimatize to the bioassay conditions and ensure no positional bias occurred regarding feeding tube preference. No positional bias was observed in any of the bioassays conducted after the initial 24 hours (*P* > 0.05). The treatment period commenced with the Eppendorf tubes being replaced with either a tube filled with 40% w/w sucrose solution spiked with treatment solution or sucrose alone (control). The bee feeding from each tube was quantified by weighing the tubes before and after each bioassay. For all laboratory bioassay experiments, see Table 1. Data was analyzed by either a Wilcoxon signed rank test or paired t-test depending on the outcomes of assumptions testing (Shapiro-Wilk) using SPSS version 26 (IBM Corp., Armonk, N.Y., USA).

**Table 1.** Laboratory bioassays conducted.

| Experiment | Choice 1 | Choice 2 |
| --- | --- | --- |
| 1 | Sucrose solution | Sucrose solution + LS |
| 2 | Sucrose solution | Sucrose solution + MS |
| 3 | Sucrose solution | Sucrose solution + LF |
| 4 | Sucrose solution | Sucrose solution + MF |
| 5 | Sucrose solution | Sucrose solution + H |
| 6 | Sucrose solution + L<sub>F</sub> | Sucrose solution + H |
| 7 | Sucrose solution | Sucrose solution + β-ocimene (**1**) |
| 8 | Sucrose solution | Sucrose solution + sabinene (**2**) |
| 9 | Sucrose solution | Sucrose solution + carotol (**3**) |
| 10 | Sucrose solution | Sucrose solution + daucol (**4**) |
| 11 | Sucrose solution | Sucrose solution + α-selinene (**5**) |
| 12 | Sucrose solution | Sucrose solution + β-selinene (**6**) |

### Analysis of carbohydrates

A modified protocol from Reiter *et al* (2018) was followed with all details provided in **Methods S1**. Five replicates per line were analysed. GC-MS data were transformed to .cdf or .mzML files and processed (ADAP chromatogram builder, chromatogram deconvolution, multivariate curve resolution) and aligned (ADAP aligner) with MZ Mine 2 (v 2.53) (Pluskal *et al*., 2010). Differences in the amount of monosaccharides (incl. fructose and glucose), disaccharides (incl. sucrose), and total sugar between lines were tested. Following tests for data normality (Shapiro-Wilk) and equality of variances (Levene’s Test), data were either analysed using an ANOVA or a Kruskal-Wallis rank sum test. Since no absolute quantification was conducted, response factors for the various sugars were not corrected for in the analysis.

### Analysis of nectar volatiles

As carrot flowers vary in size and weight, we decided not to focus on the absolute amounts of compounds (although, as sampled, the two sterile lines contained significantly less amounts of total volatiles compared to the male fertile lines LF and MF, and the open pollinated (OP) line H), but instead focussed on the differences in relative amounts between the lines. Due to the low nectar volumes in carrot flowers there is no available method to separate the nectar from the remaining floral tissue without cross-contamination, making it practically impossible to treat nectar as a separate entity in the analyses. In nectar-rich flowers, extraction with micro-capillaries, or other physical separation methods, can be used (Morrant *et al*., 2009), while in nectar-poor flowers like carrot, we are limited to centrifugation, which likely will extract some sap with the nectar, or dipping, which will extract any compounds that are to some extent water soluble from the surface of the flowers.

The floral volatile extracts in dichloromethane were analysed with the same instrumental setup and method as for the carbohydrate analysis, but without derivatization. Five replicates per line were analysed. Two samples were excluded from the dataset due to failed extractions.

### Analysis of minerals

Each nectar sample was accurately weighed and digested in 100 µL of concentrated nitric acid. The volume was made up to 5 mL with Milli-Q® purified water. Five replicates per line were run. A blank was prepared in the same way. Standards for Ca, K, Mg, Na and Sr were prepared from 1000 ppm standards (HPS, North Charleston, USA). An Agilent Technologies 5100 ICP-OES was used for the analysis, set to measure line intensities in axial mode to increase sensitivity. An ionisation suppressant consisting of 0.5% w/v CsCl was used. The following emission lines were measured: Ca: 393.366, 396.847, 422.673 nm; K: 766.491, 769.897 nm; Mg: 279.553, 280.270 nm; Na:588.995, 589.592 nm and Sr 407.771, 421.552 nm.Data were checked for normality (Shapiro-Wilk normality test) and equality of variances (Levene’s test). To test for differences between line either an ANOVA followed by a pairwise T-test, or a Kruskal-Wallis rank sum test followed by a Mann-Whitney U-test were conducted in R v 3.4.2.

### Floral semiochemical analysis and identification

Individual flowers were removed from umbellets by scissors. Flowers from 3 umbellets were used per sample. The flowers were extracted with dichloromethane (500 µL) for 24 hours in 2-mL vials, before the extracts were individually transferred to new vials. Five replicates per carrot line were analysed. Gas chromatography-mass spectrometry (GC-MS) analysis was performed as for the carbohydrate analysis, but injections (1 µL) were performed in splitless mode (1 min). Differences in the total floral volatile amounts between lines were tested for in two analyses: testing between all five separate lines, and testing between three groups of lines. For this second analysis, lines were pooled into three groups: Low yielding lines (LS and LF), medium yielding lines (MS and MF) and OP, high yielding line (H). Following tests for data normality (Shapiro-Wilk) and equality of variances (Levene’s Test), data were either analysed using an ANOVA followed by pairwise t-tests, or a Kruskal-Wallis rank sum test followed by pairwise Mann-Whitney U tests where appropriate. A Holm correction was used in the pairwise analyses. As the GC-MS data contained zero values, data were fourth root transformed, centered, and scaled prior to multivariate analysis (Wold *et al*., 2001; Hervé *et al*, 2018). To visualize differences between samples, a principle coordinate analysis was generated from a Euclidean distance matrix, using the packages ‘vegan’ (Oksanen *et al*, 2018) and ‘ape’ (88) in R v 3.5.1 (R Core Team, 2018). Candidate compounds were tentatively identified by comparisons to mass spectral databases (Wiley 9, NIST17) and were purchased or isolated from commercially available essential oils, for details please see **Methods S1**.

## Results

In the first part of the study, which consisted of a set of preliminary qualitative experiments, the responses of free-flying honey bees to carrot nectar extracted from the four parental carrot accessions (2 reciprocal pairs) were investigated in field-based bioassays. The four accessions were selected based on available seed yield data (Figure 1A, data provided by Rijk Zwaan Pty Ltd, Australia). Two reciprocal pairs of male-sterile CMS and fertile maintainer accessions were chosen as representative low-yielding (LS = low yielding sterile and LF = low yielding fertile) and medium-yielding (MS = medium yielding sterile and MF = medium yielding fertile) pairs. To determine the attractiveness of the nectars to honey bees, nectar of flowers from each parent line were extracted and presented in feeders located in the vicinity of two full size honey bee hives.

These nectars failed to elicit any honey bee attraction even after several hours. To control for sugar content (and hence reward) between the samples, we next developed a set of experiments where equal volumes of carrot nectar were added to aqueous solutions of honey containing 40% sucrose. These solutions were assessed for honey bee attraction utilising a dual-choice design with choices of either honey solution spiked with carrot nectar (treatment) or honey solution alone (control). In these experiments, with all four accessions, ≤ 2 bees fed on treatments, while on average ˃15 bees fed on controls. Two extraction methods, spinning (spun) and extraction with water (dipped), yielded comparable results. (Figure 1B) These field-based experiments indicated that all four parental carrot accessions likely contained components unattractive to bees, as the bees approached but seldom landed on the treatment feeders, but frequently landed and fed from the controls. Furthermore, those that did land on the treatments hesitated to taste and never consumed all of the solution within the feeders whereas those that landed on the control feeders consumed all (50 µL) of the honey solution. Guided by the field bioassay observations showing that carrot nectar deterred approaching bees from both landing and feeding on feeders, we designed a laboratory-based experiment combining both odour and taste factors. To facilitate quantitative experiments, a dual-choice bioassay in small cages was developed. Furthermore, a fifth high-yielding, OP cultivar, known to be relatively attractive to honey bees, Western Red (H = high yielding), was included in the lab bioassay experiments. To rule out that no other factors of the honey solutions affected the feeding, plain sucrose solutions were used as controls instead of honey solutions in these experiments. Again, addition of carrot nectar from any of the four parental lines to sucrose solutions significantly reduced feeding (P < 0.001, n = 10, Figure 1C). The amount of nectar consumed by the bees showed a trend (r(3) = .85, *P* = 0.071) that corresponded to the differences in the reported seed yield for each line (i.e. bees consumed more nectar from lines that had higher seed yields). In a more detailed dual-choice experiment, sucrose solutions were spiked with nectar from the high yielding OP H compared with sucrose solutions spiked with nectar from parent accession LF (low yielding fertile line), bees consumed significantly less nectar from line LF than from H (*P* = 0.031, Figure 1D). Based on these observations, nectar from the four parental accessions (LS, MS, LF, MF) and the OP H cultivar were subjected to detailed chemical analysis to determine which chemical factors contributed to the lack of both initial attraction and continuous nectar feeding. Odorous repellents as well as minerals were targeted in the search for anti-feedants. To determine any differences in potential nectar rewards, the levels of simple monosaccharides and disaccharides (fructose, glucose and sucrose) in the nectar extracts from parent accessions LS, MS, LF, MF and OP H were analysed. Although slightly higher sugar concentrations were observed in H, no significant differences in the amount of monosaccharides, disaccharides, or overall total sugars were found between the accessions (*P* > 0.05, Figure 2A).

**Figure 2.**
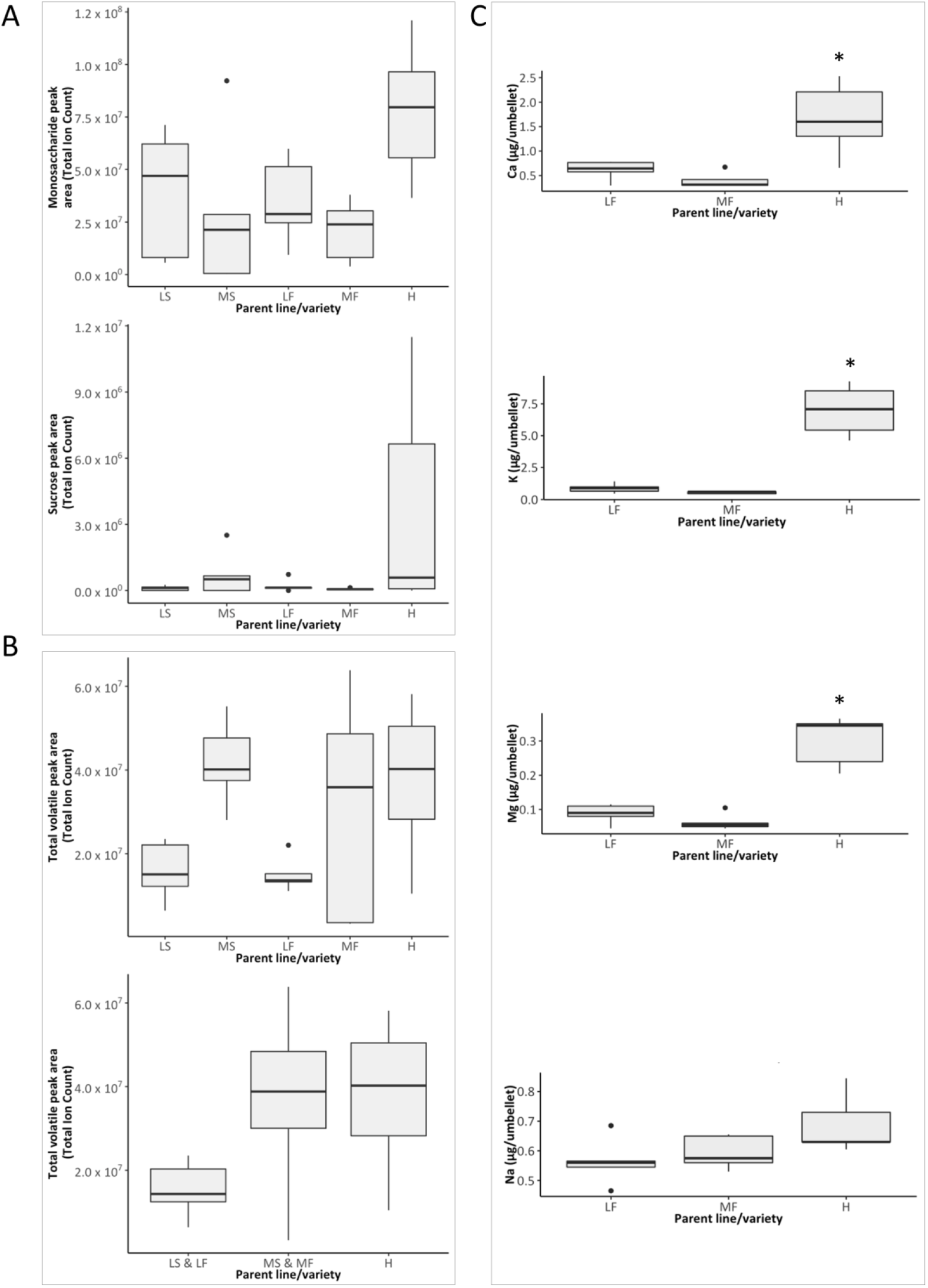
A: Relative amounts of monosaccharides and sucrose in the sampled carrot accessions. B: Relative amount of total volatiles a) in the five carrot accessions, b) in each of the three groups: low yielding pair (LS and LF), medium yielding pair (MS & MF) and high yielding open-pollinated variety H. C: Content of Ca, K, Mg and Na in parent lines LF and MF, and variety W (male sterile lines not analysed). Boxes indicate interquartile ranges with the inner line denoting the median value.

Atomic Emission Spectroscopy (AES) analysis of nectar from all fertile lines (LF, MF and H) found that the H variety had a significantly greater amount of K and Mg than did either fertile parents LF and MF (Mann Whitney U-tests, *P* = 0.024 for all), and significantly greater amount of Ca then MF (Mann Whitney U-test, *P* = 0.048). No significant difference in Na was observed (ANOVA, *P* = 0.07; Figure 2C). Male sterile lines were not analysed.

To determine whether the nectars contained odorous repellents, we undertook GC-MS analysis of the organic profile of the aqueous nectar extracts, by extracting the aqueous portion with an intermediately polar organic solvent, dichloromethane. Analysis of the organic extract indicated clear differences between the lines though the extremely low concentrations of volatiles in carrot nectars made it difficult to quantify these differences. However, as has been reported for other plant species (Raguso, 2004), organic solvent extracts of whole flowers showed the same compounds were present in much higher quantities in our carrot floral samples, allowing multivariate quantitative analysis and comparisons between lines to pinpoint the differences in attraction. We used this methodology to assess 23 samples with 112 unique compounds detected.

Before investigating any specific compounds, the total amount of floral volatiles was analysed, revealing that there was no significant difference in the amounts of total volatiles between each of the five carrots lines individually (Kruskal-Wallis rank sum test, *P* = 0.09, Figure 2B). When analysed by male sterile/fertile pairs ((LS & LF) vs (MS & MF) vs H), a significant global difference in total volatile amount between these groups was observed (*P* = 0.01, ANOVA) with the low yielding (LS & LF) having a lower total volatile concentration than the medium yielding (MS & MF; *P* = 0.03, pairwise t-test) and high yielding H (*P* = 0.02, pairwise t-test, Figure 2B) carrot lines. The principal coordinate analysis of the content of volatiles showed the five carrot accessions to have broadly overlapping clusters of compounds (Figure 3). Separation was observed along both the first and second axes. Fertile accessions LF & MF separated from male sterile LS and OP cultivar H along the first axis, with male sterile line MS occurring in the middle. Line pair LS & LF separated from pair MS & MF along the second axis, with OP H occurring in the middle of these two groups. Cumulatively, the first two axes contributed 38.4 % of the total variation (Axis 1: 23.1%, Axis 2: 15.3%).

**Figure 3.**
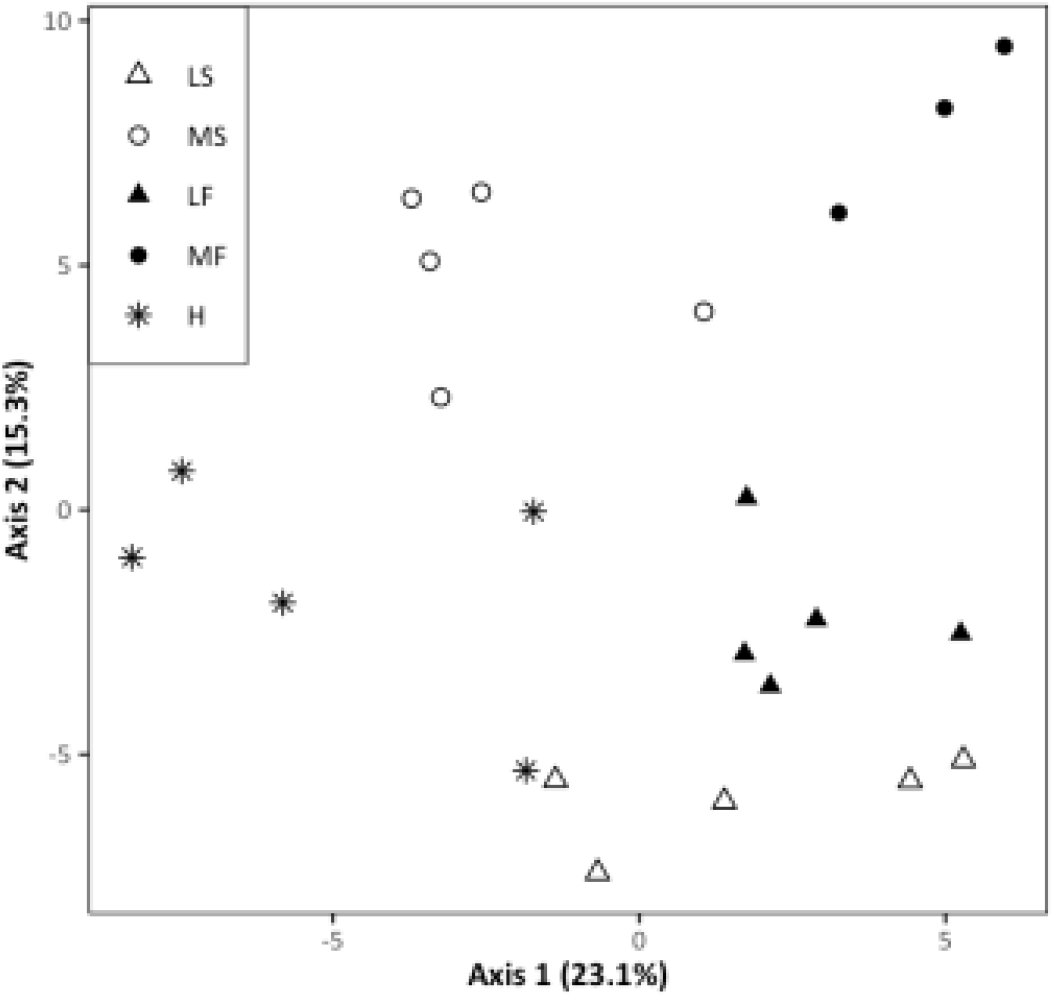
Principal coordinate analysis based on the abundance of 112 compounds detected in extracts from carrot accessions (LS, MS, LF, MF) and cultivar H. The relative corrected Eigen values denoting the percentage contribution of each axis to the total variation is displayed in the axes titles.

Compounds that showed obvious differences in abundance by GC-MS analysis between or among any accessions, and could be reliably identified, were further investigated (Figure 4A). Primarily, compounds most abundant in the accessions with lowest recorded seed set were targeted as these compounds are more likely to be repellent. The first comparisons among the parental accessions revealing a distinct difference between the male sterile and fertile pairs in the LS & LF pair, in which the monoterpene sabinene (**2**) was more abundant in the male-sterile accession (LS) (*P* = 0.007, Mann Whitney U test, Figure 4B), and was therefore considered a candidate honey bee repellent. Furthermore, the sesquiterpene alcohol daucol (**4**) was identified as a potential attractant compound, as it was found to be less abundant in the low yielding pair (LS & LF) than in the medium yielding pair (MS & MF, *P* = 0.0044, Mann Whitney U test). In the comparison between all accessions, including OP H, two sesquiterpenes; α-selinene (**5**) and β-selinene (**6**), were identified as potential repellent candidates, as these compounds were found to be more abundant in the lowest yielding lines (LS & LF) (Figure 4A) than in the medium yielding lines (MS & MF, *P* < 0.001, *P*= 0.003, respectively), Mann-Whitney U test). Additionally, the monoterpene β-ocimene (**1**) was identified as a candidate attractant, as it was found to be less abundant in the low yielding lines (LS & LF) than in the medium yielding lines (MS & MF, *P* < 0.001, Mann Whitney U test) and OP H variety (*P* = 0.002, Mann Whitney U test, Figure 4A).

**Figure 4.**
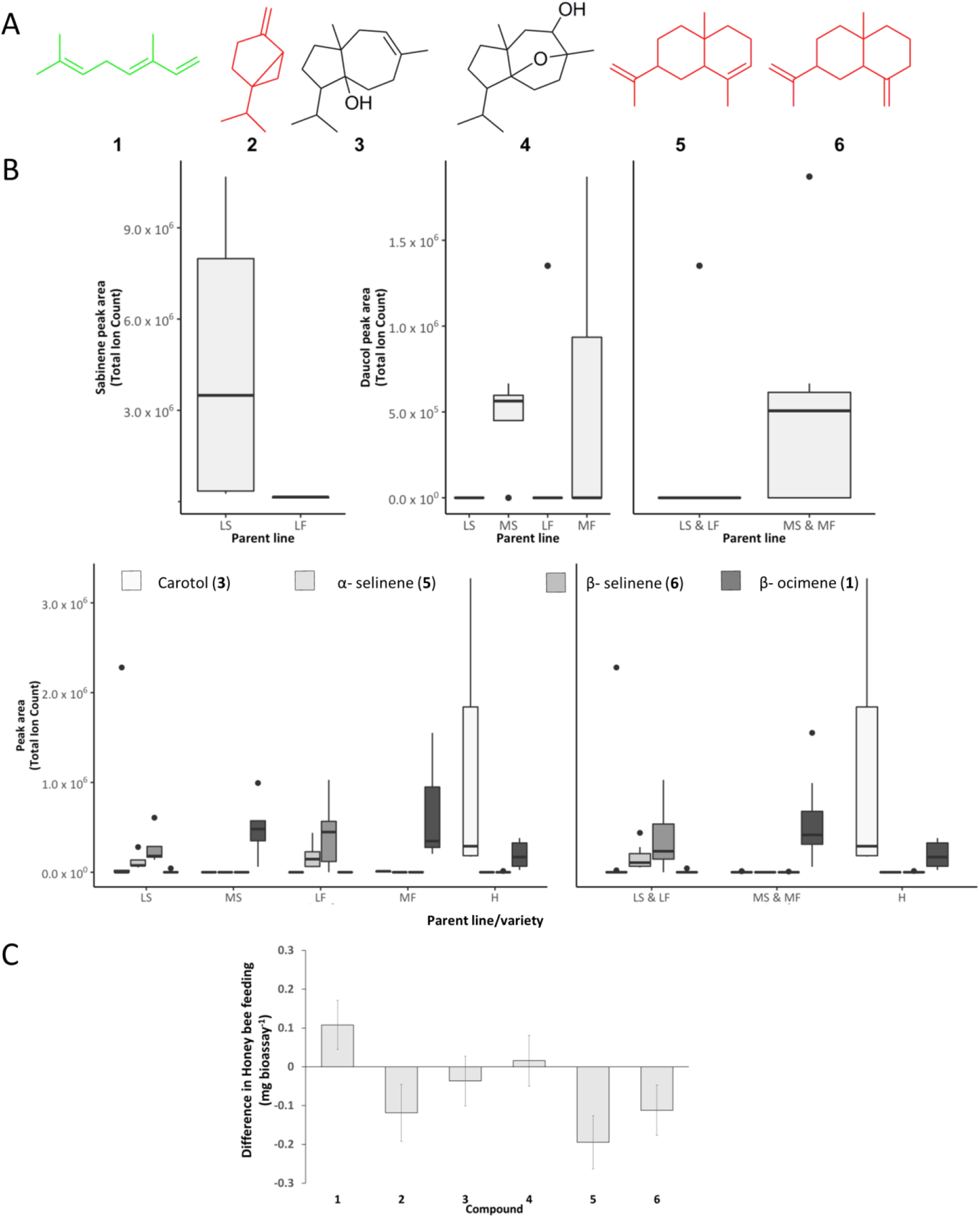
A Compounds identified from floral extracts and nectar extracts from carrot accessions LS, MS, LF, MF and H: **1** = β-ocimene, **2** = sabinene, **3** = carotol, **4** = daucol, **5** = α-selinene, **6** = β-selinene. Structures in red indicate reduced feeding, black neutral, and green increased feeding by honey bees when the respective compound was added to sucrose solutions in choice tests. B) Occurrence of identified compounds in the carrot accessions (LS, MS, LF, MF and H). C: Laboratory bioassay results of pairwise choice trials feeding of sucrose solution vs sucrose solutions spiked with synthetic compounds. Boxes indicate interquartile ranges with the inner line denoting the median value.

Carotol (**3**), was also identified as a potential attractant compound due to its greater abundance in the high yielding OP H than in both the low yielding (*P* = 0.013, Mann-Whitney U test) and medium yielding parent lines (*P* = 0.008, Mann-Whitney U test) (Figure 4B). All candidate compounds were purchased or isolated from commercially available essential oils (identity confirmed by NMR-analysis, and co-injections with extracts using GC-MS).

Candidate repellent and attractant compounds were tested in dual-choice laboratory bioassays, where bees could again choose to feed from a pure sucrose solution (control), or a sucrose solution spiked with a candidate compound (0.1 mg, treatment). After a 24-hour period, bees had consumed a greater volume of control sucrose solution compared to the sucrose solution spiked with the candidate repellents α-selinene (P = 0.012) (**5**), β-selinene (P = 0.018) (**6**), and sabinene (**2**), although this latter difference was not found to be significant (P = 0.06, Figure 4C). For β-ocimene (**1**), a candidate attractant, the bees consumed more of the spiked sucrose solution than they did of the control sucrose solution (P = 0.029), while for carotol (**3**) and daucol (**4**) there was no significant difference in the amount of the sucrose solutions consumed (P = 0.307 and P = 0.98 respectively).

## Discussion

In this study, we used a comprehensive approach to identify specific traits affecting honey bee attraction to parental accessions and open-pollinated carrot flowers. Considering the need for cross-pollination, only traits present in both male-sterile and fertile parent accessions, such as nectar and floral volatiles (but not pollen), were targeted (Gaffney *et al*., 2020). The results from our initial field bioassays suggested that the carrot nectars were not only lacking in attraction but were in fact repelling bees from the feeders. These observations guided us to develop a new laboratory bioassay protocol, based on the concept by Kessler, *et al*. (2015), in which we were able to evaluate the combined effect of odour and taste. Results from our initial laboratory bioassays with nectar from each parental accessions, revealed a trend between the relative lack of honey bee attraction in the trials, and the recorded low seed yield per line. Further, there was a significant difference between the amount of nectar that bees consumed from the low yielding fertile parental accession LF (low consumption) and the high yielding open-pollinated Western Red (H, high consumption) in these bioassays, again matching the known honey bee attraction to these lines. Chemical analyses of various plant types have demonstrated that nectars contain many more types of compounds than sugars and nutrients that provide the pollinators with a floral reward. Indeed, nectars contain a suite of chemical compounds that can act as anti-feedants, which can even be toxic to pollinators as shown in many studies, for example in ant- and honey bee pollination (Feinsinger & Swarm, 1978; Majak *et al*., 1980; Hagler & Buchmann, 1993). Secondary metabolites such as some phenolic compounds, iridoids and alkaloids (Stephenson, 1982; Kessler & Baldwin, 2007; Stevenson *et al*., 2017), carbohydrates such as xylose (Allsopp & Jackson, 1998), or inorganic elements such as potassium (Waller et al., 1972), have been shown to deter pollinator visitation. Why secondary metabolites, including anti-feedants and repellents, are incorporated in plant nectars is largely unknown (Stevenson *et al*., 2017) and references within). In many plants there is a strong correlation between the secondary metabolites in nectar and in other floral tissues, with the same compounds often present in nectar, but in lower quantities (Stevenson *et al*., 2017).

In agreement with previous studies (Gaffney *et al*., 2020), we show that the sugar levels and composition in our samples varied greatly between individuals, but no significant difference between accessions was observed. Our results show that sugars, and thereby reward quality was not linked to the observed differences in bee attraction between the accessions. In our Atomic Emission Spectroscopy (AES) analysis of minerals we showed that the concentration of calcium, magnesium and potassium differed significantly between the parental accessions and the open-pollinated Western Red (H) cultivar, despite being grown in the same soil. Notwithstanding these differences, the mineral contents were all more than ten times lower than the levels previously reported as biologically significant in nectar from other crops, such as potassium levels in non-attractive onion (Hagler, 1990) and avocado (Afik *et al*., 2006) nectars. Furthermore, the most attractive accession (H), contained the highest amount of the putatively repellent minerals (e.g. potassium), suggesting these do not constitute a major factor of the observed weak attraction to parental carrot nectars. For volatiles, we identified the same characteristic compounds both in extracts of nectar and in whole floral tissues. All identified discriminatory compounds were terpenes or terpene alcohols, all of which are relatively non-polar in nature. Despite the low polarity, these compounds were isolated from aqueous nectar solutions, which led to the design of a bioassay where a small amount of each semiochemical could be incorporated into the aqueous sucrose solutions. Taking advantage of the partial water-solubility of our secondary metabolites, the emulsion of an organic compound in an aqueous solution would represent a mimic of the natural flower, providing a slow release of odours (smell) while the bees are also exposed to the test compounds within the sugar solution (taste). Furthermore, this study relied on access to carrot accessions growing under the same controlled conditions. By collecting flowers at the same point of development, we benefitted from being able to analyse highly homogenous samples, suitable for GC-MS analysis and multivariate data treatment, allowing reliable identification of candidate bioactive compounds. The clear correlation between the presence or absence of compounds, selective feeding by bees naïve to carrot in our lab bioassays and documented seed yield for each accession, indicate that we developed an effective bioassay allowing the identification of several confirmed repellent and/or anti-feedant compounds from our samples.

Monoterpenes and sesquiterpenes, such as those identified in this study, are already known from previous pollination studies. β-Ocimene (**1**) has been suggested to be a pollinator attractant (Farré-Armengol *et al*., 2017), and has been reported as a main constituent of the bouquet of floral volatiles emitted by wild parsnip (*Pastinaca sativa*), a basal relative of cultivated carrot (Zangerl & Berenbaum, 2009). Remarkably, despite the many suggested roles for β-ocimene in biological systems (Farré-Armengol *et al*., 2017), our study is the first to confirm this compound to be an attractant to non-conditioned honey bees. It is also important to note that by adding this compound to the same sucrose solution matrix as the repellent compounds, the mixture becomes more attractive to the honey bees, serving as a control of the bioassay design. Similarly, it has been shown in a study on Asteraceae that floral odour bouquets spiked with sabinene (**2**) as part of a more complex mixture, reduced honey bee attraction (Larue *et al*., 2016). For sesquiterpenes, it has been reported that β-caryophyllene and β-elemene are attractive to *Apis cerana* (Zhang, 2018), while β-*trans*-bergamotene is believed to have an attractant effect on bumble bees (Haber *et al*., 2019). Terpenoids are also well known anti-feedants (herbivore defence) and antimicrobials (pathogen defence) (Mithöfer & Boland, 2012). Thus, from an evolutionary perspective there is likely to be a trade-off between seed set and being eaten or infected. For example, the key repellent compound found in our study, β-selinene (**6**), is a known antifungal compound in the roots of maize, and also induced by jasmonic acid in celery (Stanjek *et al*., 1997; Ding *et al*., 2017). Furthermore, previous studies on *Brassica rapa* have shown that the evolution of most floral traits are affected by insect pollination and herbivory, showcasing the importance of these interactions for plant evolution (Ramos & Schiestl, 2019). For breeding purposes, it would be fundamental to monitor and attempt to optimise the balance between pollinator attraction and plant defence. It may be suggested that the rapid anthropogenic environmental change and artificial selection in cropping systems could have disrupted this balance, which should be addressed in future breeding programs.

In conclusion, we unambiguously show that individual compounds isolated from carrot nectar and floral extracts directly impact feeding of honey bees in behavioural bioassays and subsequently impact on pollinator visitation in carrot seed production. This finding is a key step towards the development of targeted plant breeding methods for the design of hybrid carrot seed crops with improved pollinator attraction and seed yield. Plant breeding programs can now target the reduction of the levels of the identified repellent terpenes β-sabinene, α-selinene and β-selinene (Degenhardt *et al*., 2003) to improve the pollination rates of these crops for increased seed production volumes. Furthermore, our developed methodology implementing chemical phenotyping of pollination semiochemicals employing GC-MS can be applied to identify traits to be modified for improved pollination efficiency in honey bee pollinated crops with low seed yields generally.

## Data availability

Data available within the article or its supplementary materials. Any additional primary data available on request from the authors

## Acknowledgments

The authors acknowledge Tim March, Arie Baelde, Alistair Gracie, Geoff Allen and Cameron Spurr for intellectual input and help with the planning of various parts of this work. BB acknowledges funding from the Australian Research Council (ARC) Discovery Early Career Researcher Award DE 160101313. The authors acknowledge the facilities, scientific and technical assistance of the Australian Microscopy & Microanalysis Research Facility at the Centre for Microscopy, Characterisation & Analysis, The University of Western Australia, a facility funded by the University, State and Commonwealth Governments.

## Author contributions

S.R.Q.and B.B designed research, all authors performed research, S.R.Q, A.M.W, G.R.F and B.B analyzed data, S.R.Q and B.B wrote first draft, all authors contributed to final paper.

