## Supplementary Information for "Critical pollination chemistry: Specific sesquiterpene floral volatiles in carrot inhibit honey bee feeding"

Contents:

S1 Field bioassay and laboratory bioassay setup

S2 NMR spectra of α-selinene and β-selinene

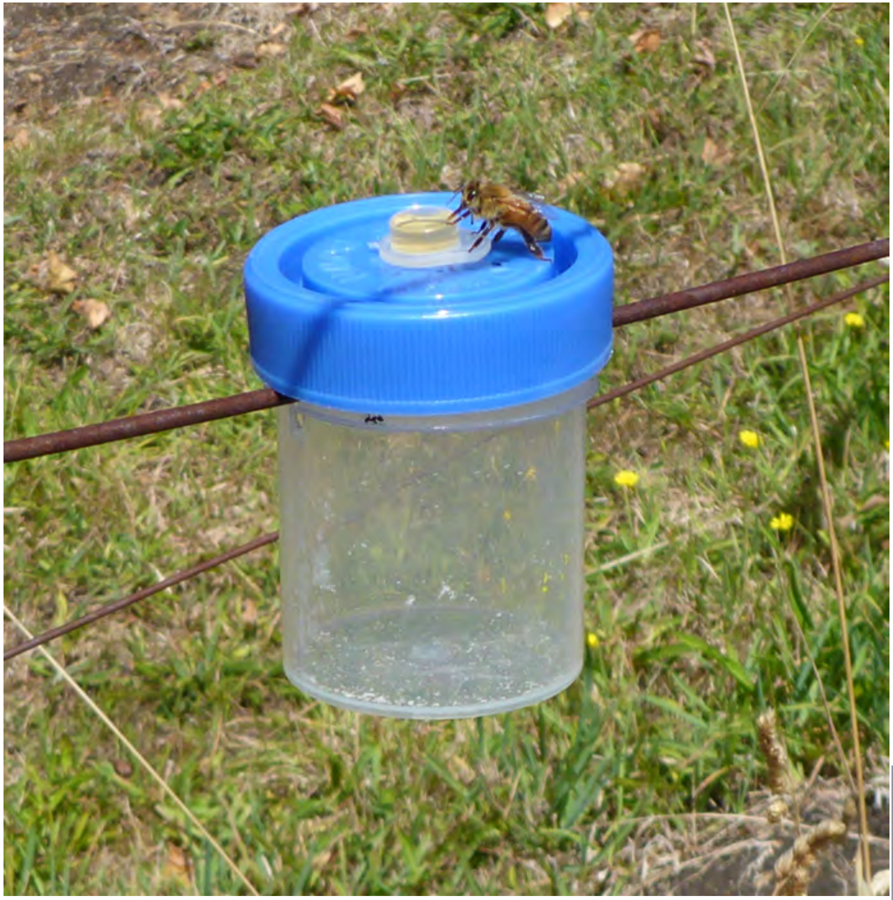

Left: Honey bee feeder
used in field bioassays

Right: Honey bee feeder
used in laboratory bioassays

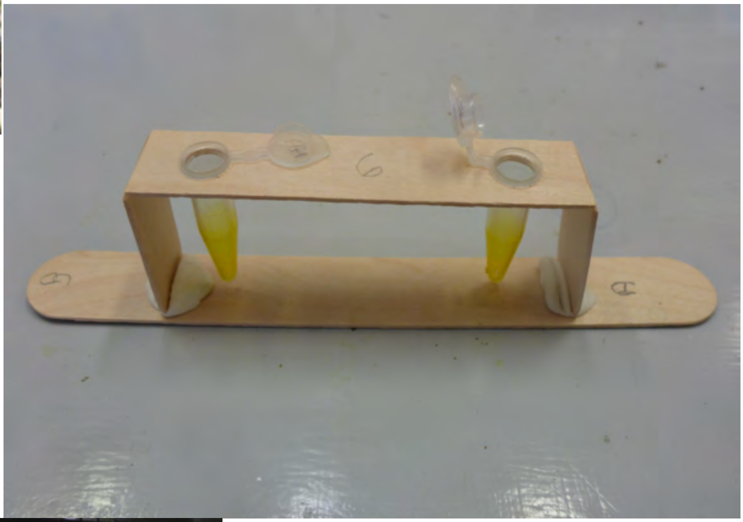

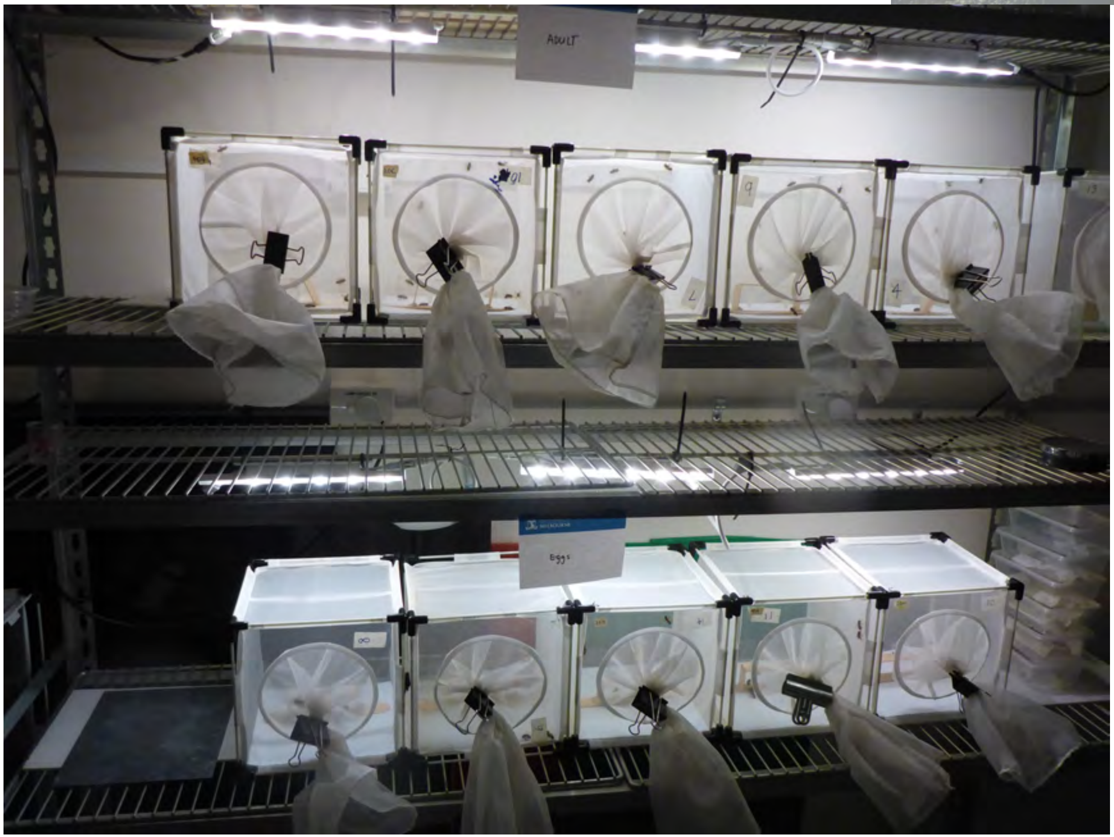


Left: Cages used for
honey bee feeder
used in field bioassays

<sup></sup>

[1]H and [13]C NMR spectra of α-selinene (5)

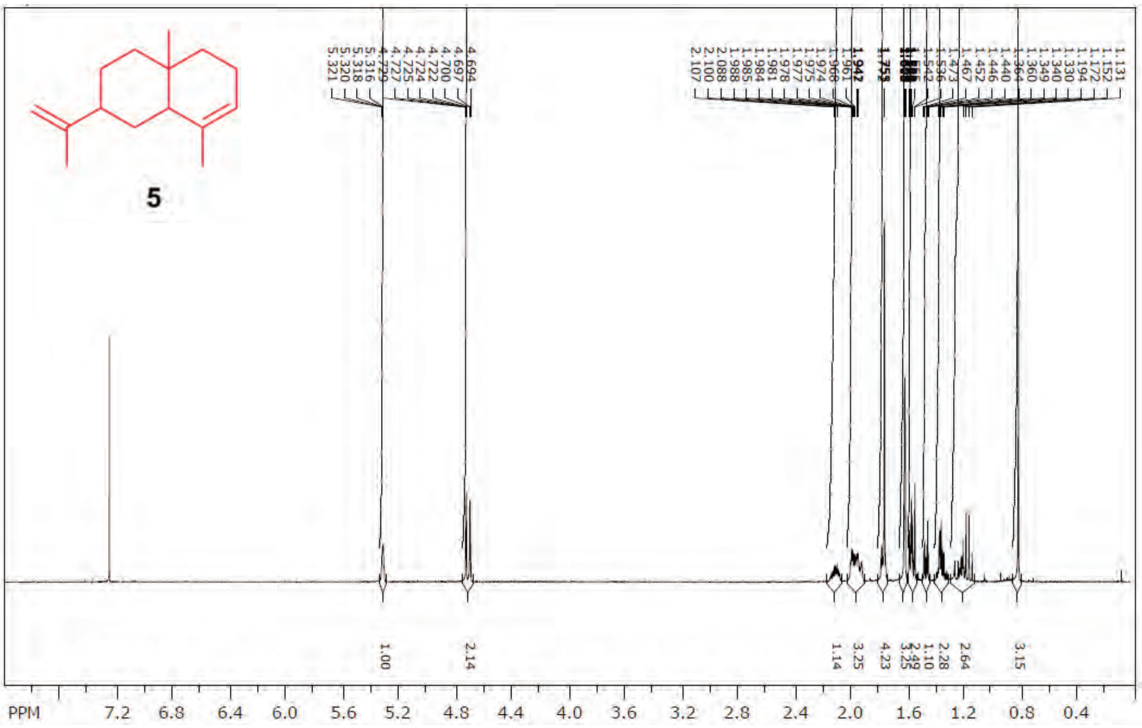

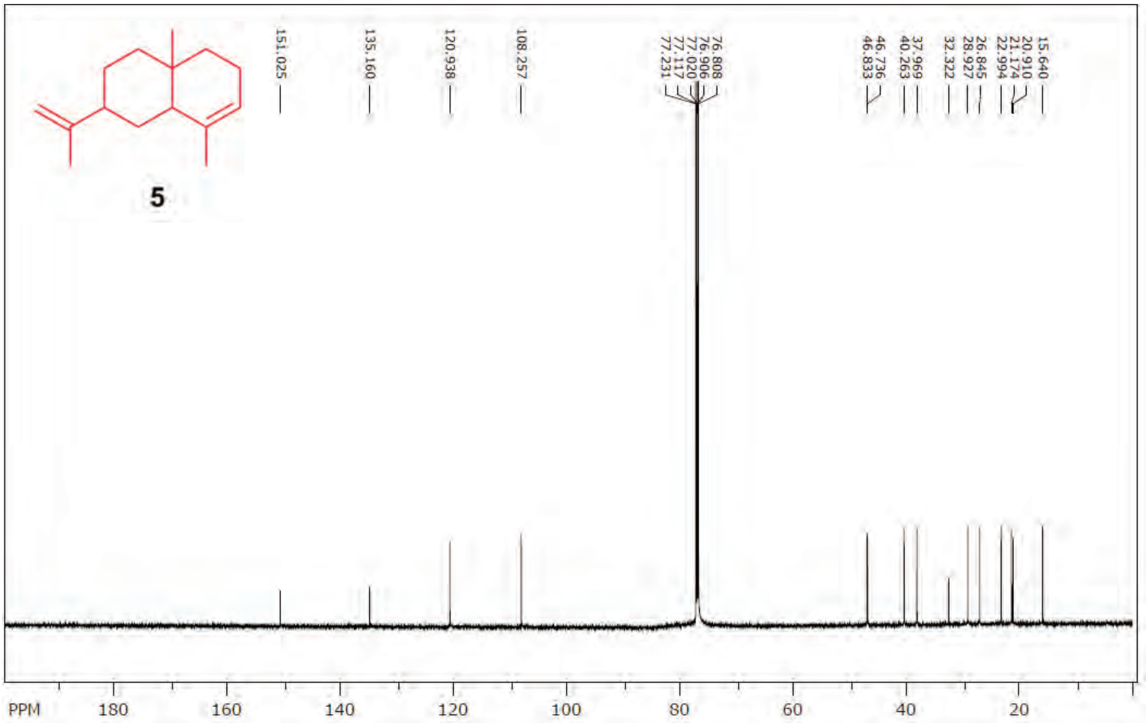

### [1]H and [13]C NMR spectra of β-selinene (6)

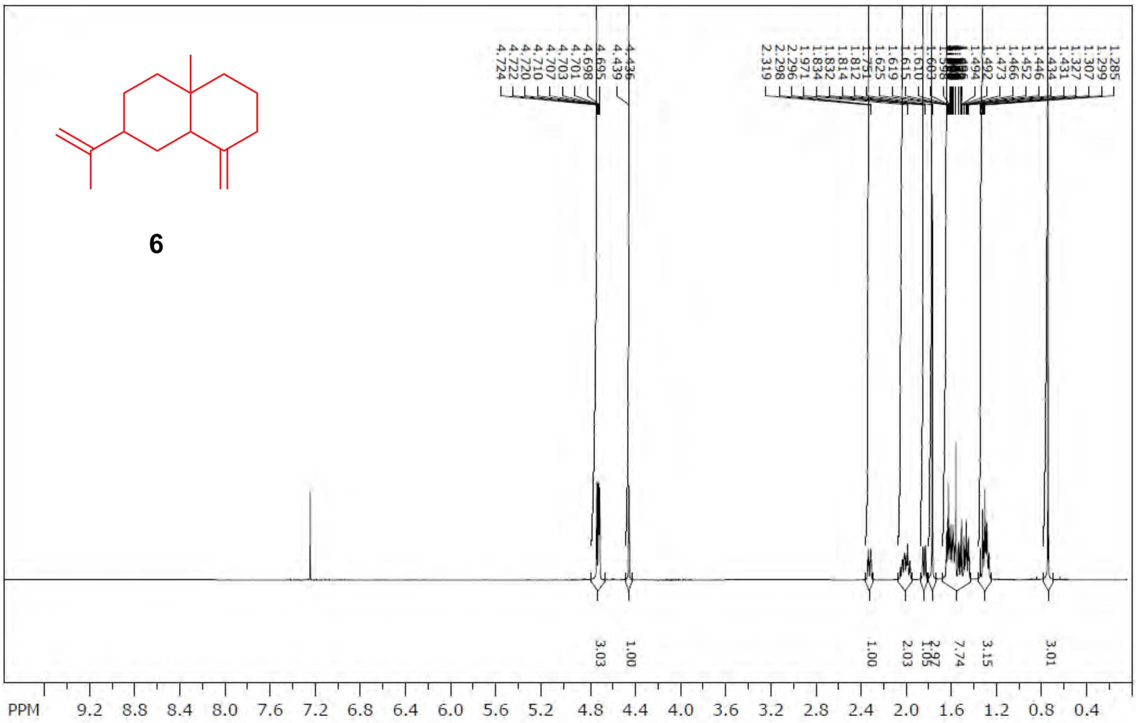

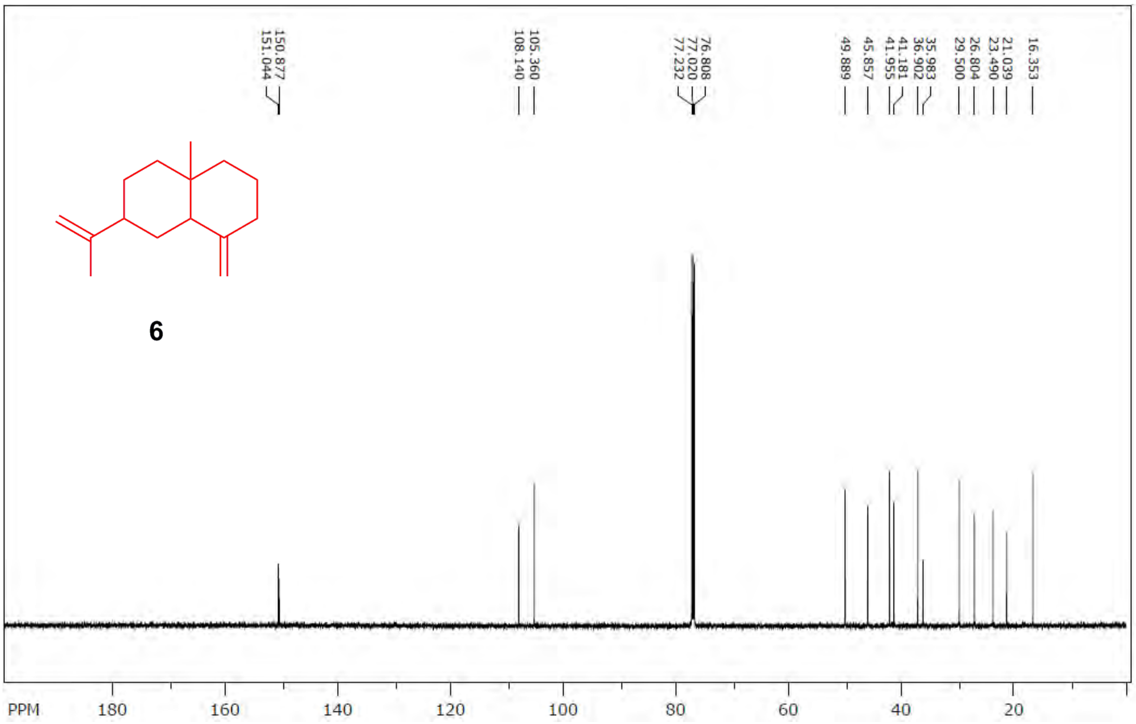
